# Continuous *cis*-regulatory changes in an advantageous gene are linked with adaptive radiation in cichlid fishes

**DOI:** 10.1101/2020.11.28.391342

**Authors:** Langyu Gu, Chenzheng Li, Xiaobing Mao, Zongfang Wei, Youkui Huang, Ximin He, Wenjun Zhou, Li Li, Deshou Wang

## Abstract

Deciphering why some lineages produce spectacular radiations while others do not provides important insights into biodiversity, but the molecular basis underlying this process remains largely unknown. Here, we identified a lineage-restricted gene, which we named *lg*. Combined omics analyses showed that *lg* is under positive selection in the most species-rich lineage of cichlid fishes, the modern haplochromine (MH) lineage, indicating its evolutionary advantage. Using transgenic zebrafish, we functionally showed that a cichlid fish-specific upstream insertion of *lg* can drive new and strong eGFP expression in tissues noted for adaptation in the MH lineage, but not in other lineages. Furthermore, the deletion of three MH-specific SNPs within this region can reconstitute weak and limited expression patterns similar to those in non-MH lineages. We thus demonstrated that a series of *cis*-regulatory changes in an advantageous gene are linked with a gain of expression that is related to an astonishingly adaptive radiative lineage.

## Introduction

Understanding how genetic changes contribute to adaptive radiation, which is characterized by closely related species diversification coupled with ecological niche adaptation in a short time, is key to understanding the emergence of biodiversity^1–3^. It is well accepted that the contribution of protein evolution to this theme is restricted due to its pleiotropic effects^4,5^. Therefore, once adaptive sequence evolution occurs, it is usually advantageous and indicates important biological functions. Indeed, such findings for proteins are often exciting but can be counted on one hand, in particular for adaptive radiation^6–8^. Instead, emphasizing the roles of gene regulation in adaptive radiation arises frequently, especially with recent advances in genome-wide association studies, in which noncoding regions are often linked with gene expression changes^9–11^. However, functional evidence of these noncoding regions in the form of their *cis*-regulatory roles, especially for small changes such as SNPs, is necessary but sparse (but see the reference^12^), particularly in the context of whether many loci with small changes or one single locus with large effects can have phenotypic changes^13^. Therefore, the molecular basis of adaptive radiation remains largely unexplored.

To investigate this question, we focused on East African cichlid fishes, a classical model of adaptive radiation^1^. East African cichlid fishes are well known for their spectacular diversity and explosive speciation, i.e., hundreds of thousands of species with closely related genomes speciated within a short time and are thus often cited as *Darwin’s dream pond*^14^ or *natural mutants*^15^. Characteristics such as anal fin eggspots, maternal mouthbrooding behaviour, skin colouration, sensory evolution and cranial cartilage together with ecological opportunities are supposed to be key factors in promoting its unparalleled species propensity^16–21^. The modern haplochromine lineage (MH) is the most species-rich cichlid fish lineage and it possesses approximately 1800 species. In contrast, most radiations of non-MH lineages of East African cichlid fishes comprise only dozens of species^22^. Although studying the adaptive radiation of cichlid fishes has been a hotspot in evolutionary biology for a long time, little focus has been placed on the genetic mechanism driving the exceptional speciation that distinguishes the MH lineage from all other cichlid fishes, except for a limited number of studies focusing on eggspot traits specific to the MH lineage^23,18^.

Combined with comparative omics analyses, computational evolutionary analyses and functional experiments, we identified a fish-specific unannotated gene, which we named *lg* here, that can be causally linked to the adaptive radiation of the MH-lineage cichlid fishes. The ancestral function of *lg* is not indispensable, considering that multiple losses occurred in different fish lineages, and no obvious phenotype was detected in gene knockout zebrafish. Interestingly, *lg* evolved an MH lineage-segregated amino acid that is under positive selection, suggesting it has newly evolved advantageous roles specific to the MH lineage. *lg* also evolved new expression profiles in the MH lineage, confirming its new functions. Indeed, *lg* evolved a cichlid fish-specific upstream insertion that can drive new eGFP expression in tissues related to adaptation, such as the pharyngeal arches and serotonergic neuroepithelial cells. Notably, this homologous region of the MH lineage and non-MH lineage drives different expression patterns, i.e., a stronger and wider range of patterns in the former. Further functional experiments showed that three MH-specific SNPs deletion can explain this difference, suggesting that small *cis*-regulatory changes can induce significant expression differences. Taken together, our work revealed for the first time that a series of *cis*-regulatory changes in an advantageous new gene are linked to an astonishingly adaptive radiative lineage.

## Results

### *lg* is a lineage-restricted gene, and its ancestral function is not indispensable

Comparative genomic analyses showed that *lg* is a lineage-restricted gene that appeared only in fishes, at least based on the available data (Supplementary Fig. 1, Table S1). *lg* was lost multiple times in different fish lineages, supporting the assumption that it is not indispensable. To further characterize its function, gene knockout was performed in zebrafish. Sanger sequencing and western blot analyses confirmed successful gene dysfunction with a frameshift (5 bp deletion) and a protein interruption (a stop codon appeared at the 18^th^ amino acid) (Supplementary Fig. 2A). Comparative transcriptomic analyses after gene knockout showed that the top differentially expressed (DE) genes were largely related to immunity, neurons and energy (Table S2). To determine whether there are different immune responses between MUT and WT, we examined the immune responses of larval zebrafish to lipopolysaccharide (LPS) by counting the numbers of *lyz*^+^ neutrophils in the caudal haematopoietic tissue one day after injection ^24^. No significant difference was detected (Supplementary Fig. 2B, Table S3). To determine whether gene knockout affects neuronal development in the brain and eyes where *lg* is expressed in zebrafish, we compared the phenotypic changes between MUT and WT using the brain neuron marker HuC ^25^ and still no difference was detected (Supplementary Fig. 2C). Then, we turned our attention to one top DE gene, *slc16a6*, which has been reported to be related to energy metabolism under fasting conditions ^26^. Under the same strategy, however, we still could not find obvious metabolic phenotypic changes between MUT and WT (Supplementary Fig. 2C). This can be explained by the fact that *slc16a6* is only downregulated here instead of entirely disrupted, as in the reference ^26^, so that it can still execute its function. These functional assays further support our assumption that the ancestral function of *lg* is not indispensable.

### *lg* is under positive selection in the MH lineage of cichlid fishes

New gene functions can evolve via protein-coding sequence evolution. We thus amplified *lg* and conducted computational evolutionary analyses to test the signal of adaptive sequence evolution in the coding region across the East African cichlid fish phylogeny (Fig. 1). Branch-site model analyses with PAML revealed that one nonsynonymous amino acid mutation (A to S) specific to the MH lineage located at the 38^th^ residue is under positive selection (*ω*>1, p<0.01) (Fig. 1, Table S4). This indicates that *lg* evolved advantageous roles in the MH lineage of cichlid fishes.

**Fig. 1.**
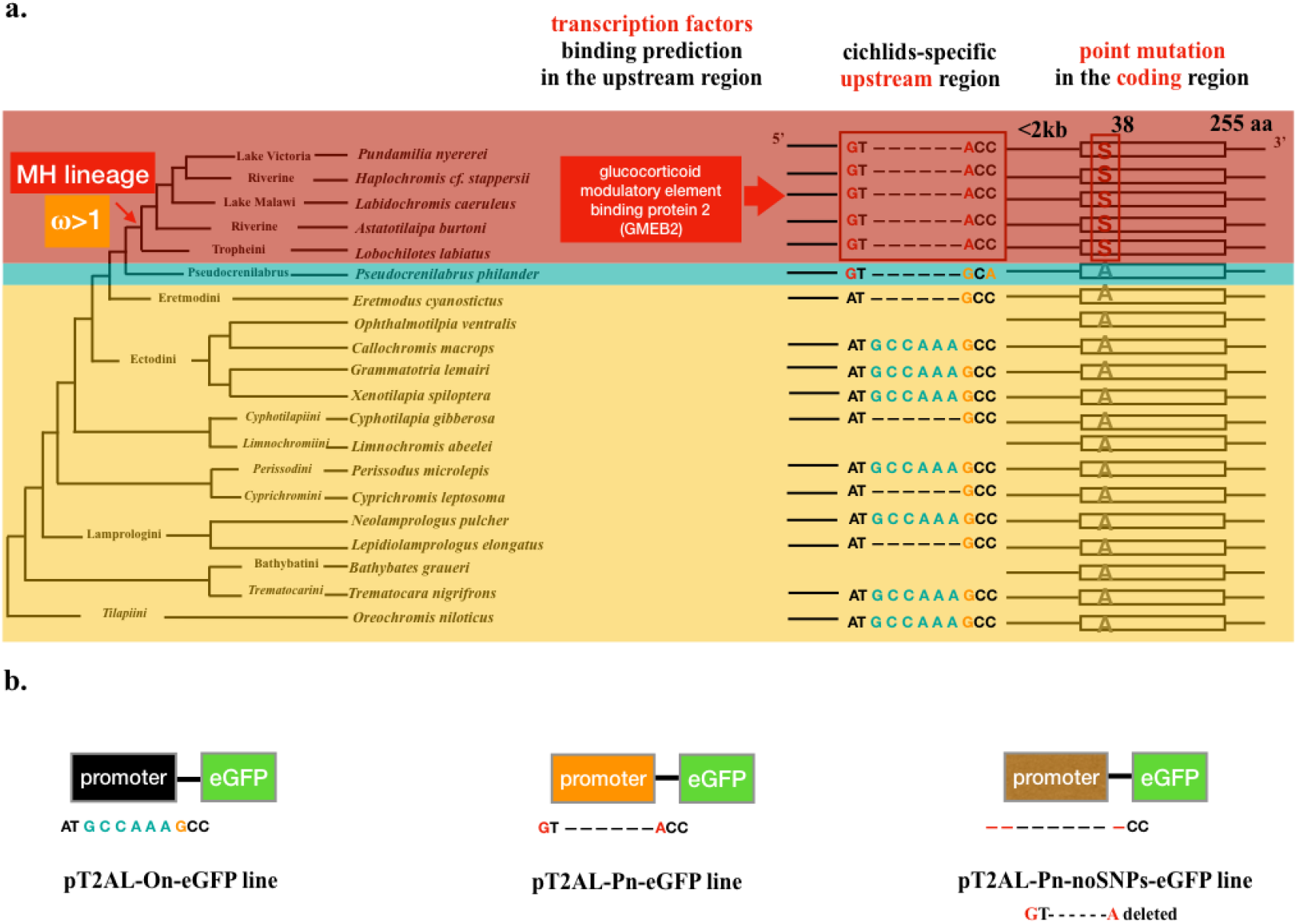
Sequence evolution of *lg* across the East African cichlid fish phylogeny. a. Both the upstream and coding regions exhibited SNPs segregated with the modern haplochromine (MH) lineage, i.e., three SNPs (GTA) in the upstream region; one nonsynonymous amino acid change (A to S) located at the 38^th^ position was under positive selection (Table S4). Transcription factor binding site prediction showed that these SNPs can set up the opportunity to bind to glucocorticoid modulatory element binding protein 2 (GMEB2). The reference tree is from the literature ^27^. b. Three transgenic plasmids were constructed. The pT2AL-On-eGFP line (On line) adopted the upstream region of *Oreochromis niloticus* as the promoter to drive eGFP expression. The pT2AL-Pn-eGFP line (Pn line) used the homologous upstream region of *Pundamilia nyererei* in the MH lineage as the promoter. The pT2AL-Pn-no-SNPs-eGFP line (Pn-noSNPs line) used the same promoter as the Pn line but we deleted the three MH-segregated SNPs (GTA).

### *lg* evolved new expression profiles in cichlid fishes, especially in the MH lineage

Changes in gene regulation can also induce phenotypic differences; thus, we examined the expression profiles of *lg* in different fishes by combining RNAseq analyses and *in situ* hybridization (Fig. 2, Table S5). The results revealed that *lg* is mainly expressed in the brain and eyes in most fishes. Intriguingly, it evolved new expression profiles in cichlid fishes in the pharyngeal cartilage, as well as in the skin in the MH lineage ^19^. These results suggest that *lg* evolved new functions in cichlid fishes, especially in the MH lineage.

**Fig. 2.**
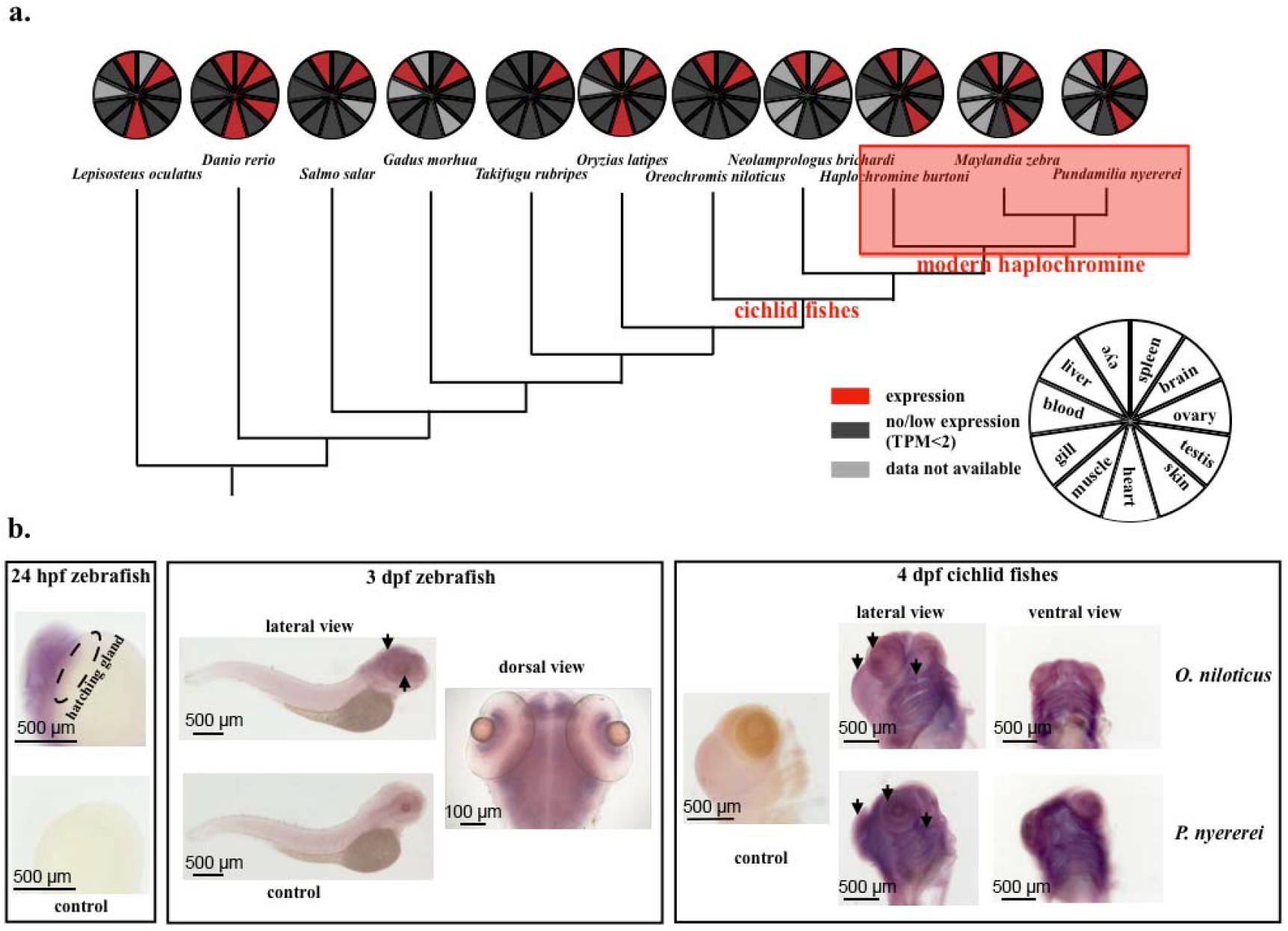
Expression profiles of *lg* in fishes. a. Transcriptomic analyses of *lg* gene expression patterns in adult fish tissues. *lg* was mainly expressed in the brain and eyes in most fishes but evolved new expression patterns in the skin in the modern haplochromine (MH) lineage of cichlid fishes. b. *In situ* hybridization in zebrafish and cichlid fish larvae. *lg* was mainly expressed in the brain, ganglion and inner plexiform cell layers in the eyes of zebrafish. No expression was detected in the hatching gland of 24 hpf (hours post fertilization) zebrafish embryos. *lg* was highly expressed in the brain, eyes and pharyngeal arches in the MH cichlid fish *Pundamilia nyererei* but weakly expressed in the non-MH cichlid fish *Oreochromis niloticus*.

### *lg* evolved a cichlid fish-specific upstream region, which possesses three MH-segregated SNPs

To determine whether there is a cichlid fish-specific regulatory region that drives the new expression patterns, we conducted comparative genomic analyses of *lg* among different teleost fishes to try to find conserved noncoding sequences (CNSs) specific to cichlid fishes (Supplementary Fig. 3). Indeed, there was one cichlid fish-specific CNS (approximately 2 kb) in the upstream region. To further characterize its sequence evolution within cichlid fishes, we amplified the whole *lg* gene, including the CNS region, in a total of 20 cichlid fishes representing main lineages across the East African cichlid fish phylogeny^27^ (Fig. 1). Interestingly, within this cichlid fish-specific CNS region, there are three SNPs specific to the MH lineage (Fig. 1). Further transcription factor (TF) binding prediction revealed that these SNPs could set up binding opportunities for a hormone-related TF, glucocorticoid modulatory element binding protein 2 (GMEB2) (Fig. 1). These results prompted us to pay special attention to the *cis*-regulatory evolution of *lg* in cichlid fishes.

### Homologous cichlid fish-specific upstream regions from the MH lineage and non-MH lineage perform different *cis*-regulatory roles

To determine whether the cichlid fish-specific upstream region of *lg* indeed has regulatory functions, and whether this region has particular function in the MH lineage, we constructed two transgenic lines with the homologous upstream regions from species belonging to different lineages as the promoters to drive eGFP expression in zebrafish (Fig. 1, Supplementary Fig. 4). One line is with the upstream region of an MH cichlid fish, *Pundamilia nyererei* (pT2AL-Pn-eGFP transgenic line, Pn line). The other line is with the corresponding homologous region of a cichlid fish belonging to a sister lineage of East African Great Lakes cichlid fishes, Nile tilapia (*Oreochromis niloticus*) ^16^ (pT2AL-On-eGFP transgenic line, On line). We successfully obtained F2 generations for both transgenic lines. Both lines can drive eGFP expression in zebrafish, confirming their regulatory functions, but with different patterns (Fig. 3 and Fig. 4). The Pn line showed strong expression in neurons in most regions of the brain (olfactory epithelium, olfactory bulb, forebrain, optic tectum, cerebellum and hindbrain), eyes, spinal cord, and heart valves. The Pn line also showed new and strong eGFP expression in the hatching gland in embryos, pharyngeal arches, chondrocyte cells in the fin and scattered epidermal serotonergic neuroepithelial cells. The positive eGFP cells were further confirmed by available antibodies (Fig. 3). Interestingly, compared to the Pn line, eGFP fluroscence was weak in the brain and pharyngeal arches in the On line (*p*<0.01), and was hard to detect in other tissues (Fig. 4, Table S6). Considering that we obtained different F1 individuals of each line possessing the same stable eGFP expression patterns, and that the eGFP expression patterns of different lines were consistent with the *in situ* hybridization and RNAseq results above, the transgenic lines we constructed were stable and credible. These results confirmed that the cichlid fish-specific upstream region of *lg* indeed evolved new regulatory functions, i.e., not only execute the ancestral expression profiles in neurons, but also evolved new expression patterns in the hatching gland and pharyngeal arches in cichlid fishes. New and strong regulatory functions further evolved in particular in the MH lineage.

**Fig. 3.**
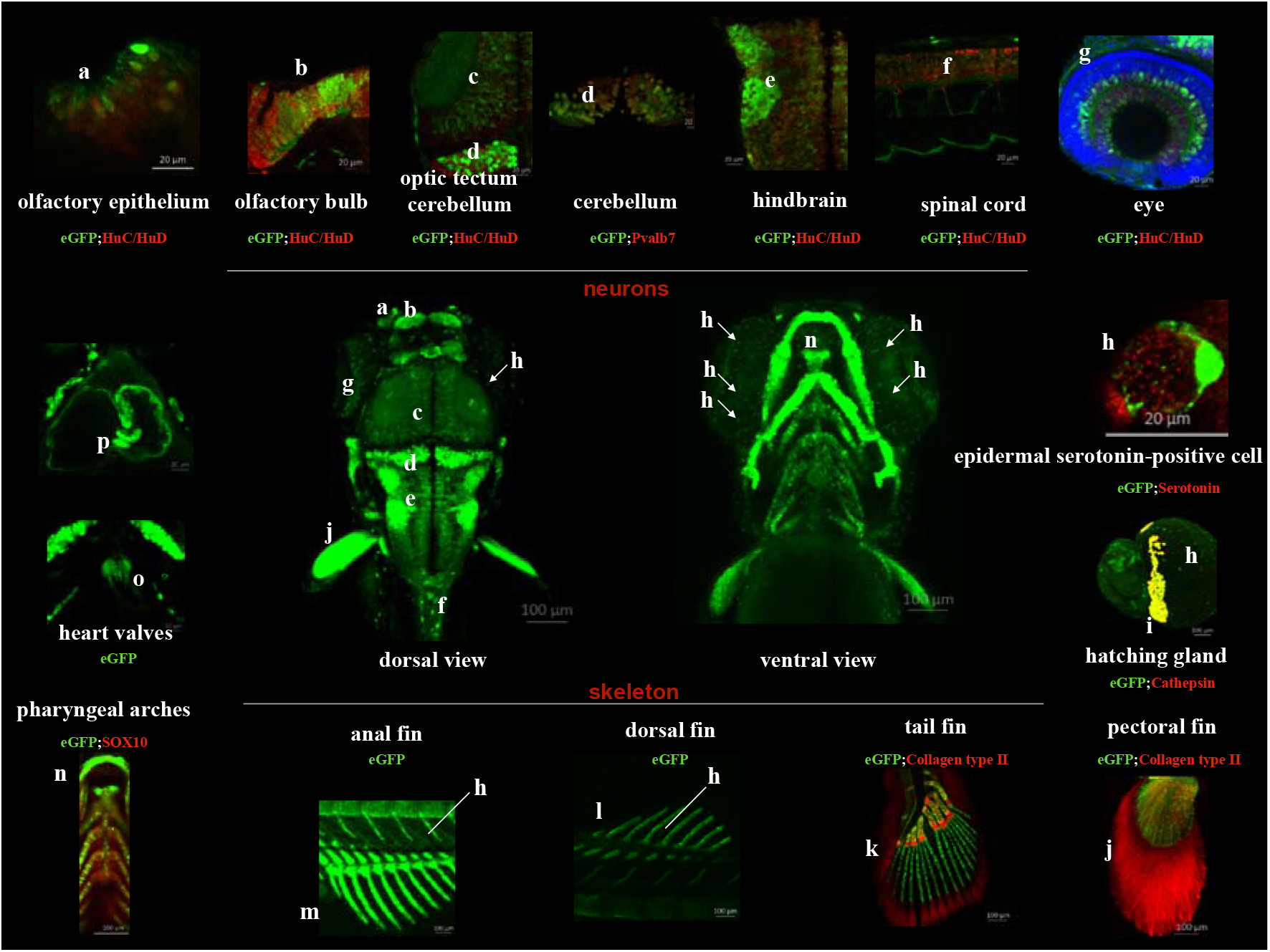
Confocal images of the expression patterns of the transgenic pT2AL-Pn-eGFP line. The green shows the fluorescent signal of eGFP. Red shows the fluorescent signal of the corresponding available cell marker antibody. Blue shows the fluorescent signal of DAPI. The merged figures are cells with double immunofluorescent staining of eGFP and the marker antibody. The pictures were taken from 4 dpf (days post fertilization) zebrafish, except the hatching gland, which is from 24 hpf (hours post fertilization) embryos. a. olfactory epithelium; b. olfactory bulb; c. optic tectum; d. cerebellum; e. hindbrain; f. spinal cord; g. eye; h. serotonergic neuroepithelial cells; i. hatching gland; j. pectoral fin; k. tail fin; l. dorsal fin; m. anal fin; n. pharyngeal arches; o and p. heart valves.

**Fig. 4.**
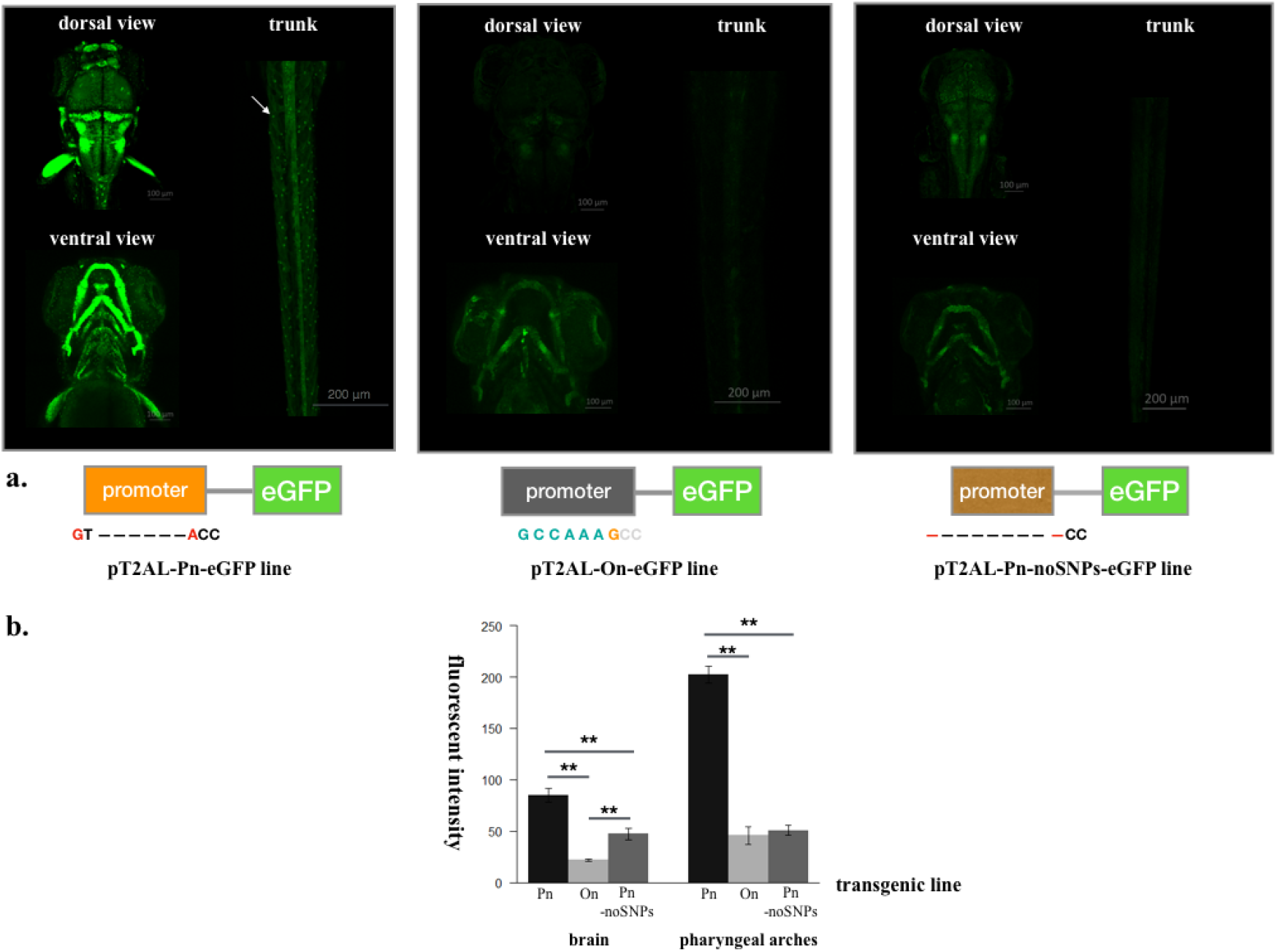
eGFP expression comparisons among three different transgenic lines. a. Different eGFP expression patterns of three transgenic lines in zebrafish. The Pn line showed strong expression in neurons in most regions of the brain, eyes, spinal cord, heart valves, pharyngeal arches, fin and epidermal serotonergic neuroepithelial cells. Compared to the Pn line, eGFP fluorescence was weak in the brain and pharyngeal arches in the On line and was hard to detect in other tissues. Only with these three SNP deletions did the Pn-noSNPs line clearly show significantly different expression patterns from the Pn line but similar patterns to the On line. b. Statistics of fluorescence strength comparisons among three transgenic lines, focusing on the brain and the first two pharyngeal arches. The Pn line showed significantly stronger fluorescence than the other two lines. With the three MH-segregated SNP deletions, the Pn-noSNP line showed weak fluorescence expression, similar to the On line. Mean and SD error bars are shown. n=6 for each measurement. Detailed statistical analyses can be found in Table S6.

### Three MH lineage-segregated SNPs are causally linked with different *cis*-regulatory roles

To determine whether the different expression patterns above were caused by the three MH lineage-segregated SNPs, we constructed another transgenic line with the same upstream region as in the Pn line as the promoter, but specifically deleted the three MH-segregated SNPs (pT2AL-Pn-noSNPs-eGFP line, Pn-noSNPs line) (Fig. 1, Supplementary Fig. 4). Interestingly, only with these three SNP deletions did the Pn-noSNPs line clearly show significantly different expression patterns from the Pn line but similar patterns to the On line (Fig. 4, and Table S6). These results confirmed that these SNPs are causally linked to the different regulatory roles of the cichlid fish-specific upstream region between MH and non-MH lineages.

## Discussion

Here, we revealed that the evolution of a lineage-restricted gene, *lg*, is causally linked with one adaptive radiative lineage of cichlid fishes, the MH lineage. Although the ancestral function of *lg* is not indispensable, adaptive sequence evolutionary analyses of *lg* discovered its newly evolved advantageous roles in MH lineage cichlid fishes. Lineage-restricted genes are increasingly being reported to be related to novelty and adaptation^28–30^. It has been shown that they are usually located at the edge of the gene network ^31^, and thus their disruption does not significantly affect the core gene network. This could be one reason why their coding regions are relatively free to evolve new functions, such as *lg* in our study. Further gene knockout in MH cichlid fishes will be helpful to unravel its protein functions and the involved pathways.

Most unique tissues that express *lg* in the MH lineage are well known to be related to adaptation. For example, the hatching gland is a key innovation in regulating the hatching period of aquatic embryos to become free-living larvae ^32^; the pharyngeal cartilages related to feeding and ecological niche exploration are key to cichlid fishes adaptive radiation ^33–35^; epidermal serotonergic neuroepithelial cells are chemoreceptors related to environmental stimuli such as hypoxia ^36^; and the olfactory epithelium is famous for the detection of odorants, such as amino acids, hormones and pheromones ^37^. *lg* is also highly expressed in the heart valves, a key innovation to control unidirectional blood flow^38^. Given that most of these tissues are derived from neural crest cells ^39^, a key vertebrate innovation*, lg* can also be causally linked to neural crest cell development and differentiation. Taken together, we can conclude that *lg* gained new regulatory functions related to adaptation in the most species-rich lineage of cichlid fishes, the MH lineage.

Notably, unlike in the MH lineage, the homologous upstream region only drives limited and weak expression patterns of *lg* in the non-MH lineage, mainly in the brain and pharyngeal cartilage. The three MH-specific SNPs are causally linked to this difference. We thus hypothesize that a series of *cis*-regulatory changes that occur in the MH lineage are responsible for driving the new expression pattern of *lg*. The insertion of upstream regions in the ancestral East African cichlid fishes provided the basic background to evolve new gene regulation roles. Under relaxed selection pressure, this region further accumulated mutations, such as the three MH-segregated SNPs in the MH lineage. This event provided new TF binding opportunities to help *lg* integrate into a new gene network and finally trigger wider and stronger expression patterns in tissues related to adaptation in the MH lineage. In this case, we provided evidence that small SNP changes can induce significant regulatory differences. Indeed, even a single mutation in the upstream region can induce new expression profiles ^12,40,41^. It will be interesting to test our predictions of the binding of a glucocorticoid hormone-related TF to these SNPs in the future. If this is true, these SNPs can efficiently connect the organism with the external environment, since glucocorticoids are well-known stress-related hormones that regulate animal behaviour in response to the environment ^42,43^. Taken together, we showed that continuous *cis*-regulatory changes linked *lg* with a gain of new and strong expression patterns in a particularly adaptive radiative lineage.

Recent comparative genomics studies on cichlid fishes have continued to find that the speciation rate is positively correlated with genetic diversity^44–46^. Genome polymorphisms can thus provide exceptional genome potential to allow for adaptive radiation. As in this case, mutations as small as SNPs accumulated in the noncoding region can immediately induce large phenotypic changes. It can thus facilitate incipient speciation rapidly due to incompatible regulation by hybridization^47^, which is particularly meaningful for adaptive radiation. The series of *cis*-regulatory changes of a new advantageous gene linked with an adaptive radiative lineage that we discovered thus provides a clue to illuminate the molecular mechanism driving adaptive radiation in a broad sense.

## Methods

### sampling

Samples of cichlid fishes for comparative genomic analyses were provided by Prof. Walter Salzburger in University of Basel, Switzerland; cichlid fish *Pundamila nyererei* for transgene plasmids construction was obtained from IGV (Interessengemeinschaft Viktoriasee-Cichliden); Strains of AB (for transgenic lines construction) and ABTü (for gene knockout) zebrafish (*Danio rerio*) were raised under standard condition (28.5°C, 14/10 h light/dark). Tilapia (*Oreochromis niloticus*) were raised at 26°C, 14/10 h light/dark. Animal experiments have been approved by the cantonal veterinary office in Basel under permit number 2317 in University of Basel, Switzerland; the Committee of Laboratory Animal Experimentation at Sun Yat-sen University and Southwest University, China.

### -omics analyses to characterize the evolution of *lg* across fish phylogeny

To characterize the sequence evolution of *lg*, tblastx was conducted with -*e* 0.001 in fishes with good genome assembly in Ensembl 100 (Table S1) and local blast v2.9.0 (tblastx) was conducted with draft genomes assembly across the phylogeny according to the reference ^48^. After obtaining hits, the scaffolds possessing the corresponding sequences were extracted to predict the coding regions with Augustus 3.3.3 ^49,50^. The predicted *lg* coding sequences were further confirmed by aligning the translated amino acids with the available sequences from Ensembl 100 using Geneious 8.0.5 ^51^.

### -omics analyses to characterize the evolution of *lg* within cichlid fishes

To further characterize the sequence evolution of *lg* within East African cichlid fishes, we amplified the whole gene sequence and its upstream region (ca.10 kb in total) in different cichlid fishes. Combined with available cichlid fish genomes, we studied the sequence evolution in a total of 20 cichlid fishes representing main lineages across the phylogeny ^27^. Genomic DNA was extracted using a DNeasy Blood & Tissue kit (Qiagen). PCR primers were designed using Primer premier 5 ^52^. High-fidelity PCR Master Mix (BioLabs) and a touch-down annealing process were used for amplification. PGM Ion Torrent was used to sequence long amplicons (400 bp) with CHIP 316 (Life Technologies). Galaxy v20.09 ^53^ was used to trim low-quality reads (window size 2, step size 1, minimum quality score 20) and filter short reads with a minimum size of 40 bp. The sequence of a cichlid fish, tilapia, retrieved from the Ensembl database was used as the reference to recheck the assembled sequences. The sequencing reads of the *lg* gene across the cichlid fish phylogeny are available at NCBI (BioProject ID PRJNA611690). The ion torrent sequencing of most species was successful except *Limnochromis abeelei, Ophthalmotilapia ventralis*, and *Bathybates graueri*, whose sequences were further obtained from Sanger sequencing (Primers can be found in Table S6).

Genomic alignment of *lg* of cichlid fishes and other teleost fishes was conducted with mVISTA ^54,55^ using the LAGAN alignment tool ^56^ with *P. nyererei* as the reference. We applied the repeat masking option with fugu as the reference. Transcription factors (TFs) binding prediction analyses were conducted with MatInspector v8.4.1 with the genomatix software suite (https://www.genomatix.de) using default parameters with zebrafish as the reference ^57,58^.

### Positive selection detection of *lg* in cichlid fishes

To detect whether positive selection occurred in the evolution of *lg* in a specific branch, a branch-site model within the codeml in PAML v3.14 was performed ^59,60^. Fixed branch lengths (fix_blength = 2) retrieved under the M0 model were used. Alignment gaps and ambiguous characters were removed (Cleandata = 1). The tree for the PAML test is based on the phylogeny from the study^27^. Rates of nonsynonymous to synonymous substitutions (*ω* or dN/dS) significantly larger than 1 indicate that the sites in a specific target branch (foreground branch) are under positive selection. A *chi*-squared approximation was calculated to test statistical significance. Detail statistics of branch-site model comparisons can be found in Table S4.

### CRISPR/Cas9 gene knockout in zebrafish

The *lg* mutant line was generated by the CRISPR/Cas9 system. The guide RNA (gRNA) target site was selected using the CRISPOR v4.98 ^61^ with a 20 bp NGG PAM motif. The synthesized gRNA and Cas9 mRNA were coinjected into one-cell stage zebrafish embryos at a concentration of 50 ng/μl. Ten randomly selected embryos at 1 dpf (days post fertilization) were lysed and used to assess functional gene knockout using PCR amplification with primers designed at the target knockout site (Table S7). Heterozygous F1 fish were obtained by crossing F0 and wild-type (WT) zebrafish and genotyped with the same primers above using Sanger sequencing at the target site (Table S7). Siblings of the F1 generation harbouring the same mutations were mated to generate homozygous F2 mutants (MUT) and WT sisters (as the control).

### Western blot

Since there was no available antibody for *lg*, we added an HA tag (ATGTACCCATACGACGTCCCAGACTACGCT) in front of the coding region of *lg* by PCR amplification (Table S6). The antibody of the HA tag can thus detect the fused *lg* protein if there is no disruption. To ligate the amplicons of HA-tagged coding regions of WT and MUT into the pCS2 vector, restriction enzyme sites (*Bam*H I and *Xba* I) were added at the two ends of the amplicons by PCR amplification (Table S7), followed by a double digestion system (NEB) and ligation (Promega) (Table S7). The plasmids pCS2, pCS2-lg-HAtag-WT and pCS2-lg-HAtag-MUT (2 μg) (Supplementary Fig. 4) were then transfected into the HEK293T cell line (ab255449, Abcam) separately. Protein was collected after 36 h of transfection to perform Western blot according to the standard procedure ^62^. Mouse anti-HA (901501 Biolegend, 1:2000) and mouse anti-GAPDH (60004-1-lg Proteintech, 1:2000) antibodies were used for detection. Detections were performed with goat anti-mouse secondary antibody (31430 Thermo 1:5000) and enhanced chemiluminescence (ECL plus) reagent.

### RNAseq analyses

To determine the gene expression profiles of *lg*, the available raw transcriptomic data of different tissues in different fishes were extracted from the SRA database (Table S5). The raw reads were mapped to the corresponding transcriptome retrieved from Ensembl using Hisat2-2.1.0 ^63^ with default parameters. The reported SAM results were converted into BAM format, and finally recorded as count files using SAMtools v1.10 ^64^ followed by TPM value calculation ^65^.

To identify the affected genes after knockout, comparative transcriptomic analyses were conducted between MUT and WT. RNA extraction was performed (each replicate contained 30 pooled individuals) using TRIzol® reagent (Invitrogen). Six biological replicates of WT and MUT of the F2 generation were used for RNAseq analyses. Library construction and sequencing were performed using Illumina TrueSeq RNAseq technology at the Novogene Company, China, with a paired end read length of 350 bp. The raw reads were mapped to the zebrafish transcriptome assembly using the same methods mentioned above to obtain the read count files. The differentially expressed (DE) transcripts between WT and MUT were further analysed using the edgeR v3.20.7 package (FDR<0.01).

### *In situ* hybridization

The gene fragments in the coding region of *lg* in zebrafish and tilapia were amplified using PrimerSTAR (Takara) (Table S7) and *lyz* probe synthesis was conducted as described in the reference ^66^. The amplified products were cloned into the pGEM®-T Easy vector (Promega). Plasmids were extracted and purified using a TIANprep Mini Plasmid Kit II (Tiangen). The insertion and direction of probes were confirmed with Sanger sequencing using M13 PCR products (Tsingke). RNA probes were synthesized with the DIG RNA labelling kit (SP6/T7) (Roche). The experiment was conducted as described in the reference ^67^ with biological replicates (n=30 for zebrafish, n=20 for tilapia, and n=10 for *P. nyererei*). The embryos or larvae were mounted in 1% low melting point agarose (GBCBIO) and imaged with a Zeiss Axio Imager Z2 microscope.

### Transgenic plasmids and transgenic lines construction

To determine the regulatory functions of cichlid fish-specific upstream regions, we constructed plasmids with different upstream regions as promoters to drive eGFP expression in zebrafish (Fig. 1 and Supplementary Fig. 4), i.e., 1) species in the MH lineage (*P. nyererei*) (Pn line); and 2) the corresponding region of species in the sister lineage of East African Great Lakes cichlid fishes (tilapia) (On line). 3) To determine whether the three MH-segregated SNPs in the cichlid fish-specific upstream region are important, we specifically deleted these SNPs using site-directed mutagenesis by PCR primer extension (Table S7) ^68^ to construct another transgenic line (Pn-noSNPs line). A schematic of the plasmid construction can be found in Supplementary Fig.4.

DNA fragments for different plasmid constructs were amplified with PCR primers supplemented with the corresponding restriction endonuclease sites (Supplementary Fig. 4, Table S7). Fragments were then recovered (Tiangen), digested (NEB) and ligated (Promega) into the pT2AL vector following the instructions accordingly. Successfully constructed recombinant plasmids were further confirmed by Sanger sequencing (Tsingke). Injections were performed into the one-cell stage of zebrafish embryos. By outcrossing with WT zebrafish, we successfully obtained F2 stable transgenic lines for all these lines (Pn line, On line and Pn-noSNPs line). The fluroscence signals were imaged with Zeiss 880 confocal microscope and Olympus FV3000 microscope.

### Immunohistochemistry (IHC)

The following primary antibodies were used: mouse anti-Pvalb7, 1:1000 provided by Prof. Masahiko Hibi, Nagoya University, Japan ^69^; rabbit anti-serotonin, 1:400 (s5545, Sigma); rabbit anti-calretinin, 1:400 (ab702, Abcam); goat anti-GFP, 1:400 (ab6673, Abcam); mouse anti-HuC/HuD, 1:1000 (16A11, Invitrogen); rabbit anti-SOX10, 1:400 (ab229331, Abcam); rabbit anti-PAX6 1:400 (ab209836, Abcam); mouse anti-collagen type II (II-II6B3, DSHB); and rabbit anti-cathepsin L/MEP (ab209708, Abcam). The secondary antibodies were donkey to goat IgG H&L (Alexa Fluor^®^ 488) (ab150133, Abcam), donkey to mouse IgG H&L (Alexa Fluor^®^ 647) (ab150107, Abcam), donkey to rabbit IgG H&L (Alexa Fluor^®^ 647) (ab150063, Abcam) and DAPI (ab228549, Abcam). Fish larvae were fixed in 4% paraformaldehyde (Sigma) for 4 h at room temperature, followed by a detailed protocol as described previously^70^. The fluroscence signals were imaged with a Zeiss 880 confocal microscope.

### Immune response detection

To determine whether there are different immune responses between MUT and WT zebrafish, we injected lipopolysaccharide (LPS), the key component of Gram-negative bacteria, into the circulation of 2 dpf zebrafish and counted the numbers of *lyz*^+^ neutrophils in the caudal haematopoietic tissue at one day post injection. The *lyz*^+^ whole-mount RNA *in situ* hybridization signals across eight somites of the larval CHT (caudal haematopoietic tissue) regions were manually counted. Statistical analyses can be found in Table S7.

## Data Availability

The publicly available datasets that this study used are provided in the supplementary information. The RNAseq data of WT and MUT are available at NCBI (BioProject ID PRJNA611476). The ion torrent sequencing reads of the *lg* gene are available at NCBI (BioProject ID PRJNA611690). The scaffolds possessing *lg* of teleost fishes across the phylogeny, the assembly of *lg* including the upstream region of cichlid fishes across the phylogeny retrieved from ion torrent sequencing and Sanger sequencing can be found in Dryad https://doi.org/10.5061/dryad.fxpnvx0qb. Source associated data of Fig. 4 are provided in the supplementary files. Other data are available upon reasonable request.

## Acknowledgements

Many thanks to Prof. Walter Salzburger, Attila Rüegg, Zhen Liu, Zhiqiang Zhou and Hesheng Xiao for the sampling and suggestions. We thank Prof. Masahiko Hibi and Prof. Kiyoshi Naruse for providing the antibody Pvalb7. We also thank Prof. Lingfei Luo, Prof. Hua Ruan, Prof. Yang Liu and Prof. Tienming Lee for the laboratory supports. Many thanks to Prof. Zuzana Musilova, Dr. Xiaogui Yi, Dr. Yandong Zhan, Dr. Fangying Zhao, Yuepeng He, Xinkai Cheng, Qing Li and Yue Gao for the supports. Thanks to Huansheng Tao and Jialing Xu for the Confocol microscope support, Zhao Long for fluroscent microscope support. We also thank Prof. Xiangjiang Zhan, Prof. Daiping Wang and Dr. Pengcheng Wang for the comments on this ms. This study was funded by grants from the National Natural Science Foundation of China (31802283); Guangdong Basic and Applied Basic Research Foundation (2020A1515010330); the Fundamental Research Funds for the Central Universities (20lgpy109); China Postdoctoral Science Foundation (2018□M633311) and Doctoral Scholarship of University of Basel, Switzerland to LG; Grants from the National Natural Science Foundation of China (31630082) to DW, and (31771623 and 31571500) to LL. Part of -omics analyses was supported by National Supercomputer Center in Guangzhou, China. The English language of this article was edited by Elsevier Language Editing Services.

## Authors contributions

LG, DW and LL conceived this work. LG carried out –omics sequencing and evolutionary analyses. LG, CL, XM and ZW conducted the *in situ* hybridization. LG carried out the IHC. XH and WZ participated the IHC experiments of serotonin-positive cells and olfactory epithelium cells. LG, CL and XM constructed plasmids of the Pn line and the On line. LG constructed plasmids of the Pn-noSNPs line. LG leads the transgenic lines and mutant lines construction with the help from CL and XM. LG took the confocal photos. YH performed the Western blot experiment. ZW carried out the LPS immune response experiments. LG conducted the fasting ORO staining experiments. LG wrote the initial manuscript draft, and all authors commented on the manuscript and approved the final version.

## Supplementary information

Table S1 *lg* appearance in species with available genomes assembly in Ensembl

Table S2 Top differentially expressed transcripts between MUT and WT zebrafish

Table S3 The numbers of LPS immune response signals comparisons between WT and MUT zebrafish

Table S4 Positive selection detection of *lg* in cichlid fishes within codeml in PAML

Table S5 RNAseq analyses of *lg* in different fishes with available data

Table S6 eGFP fluroscence strengths comparisons among different transgenic lines

Table S7 PCR primers and annealing temperatures

Supplementary Figures

