## Supplementary Figures for "Continuous *cis*-regulatory changes in an advantageous gene are linked with adaptive radiation in cichlid fishes"

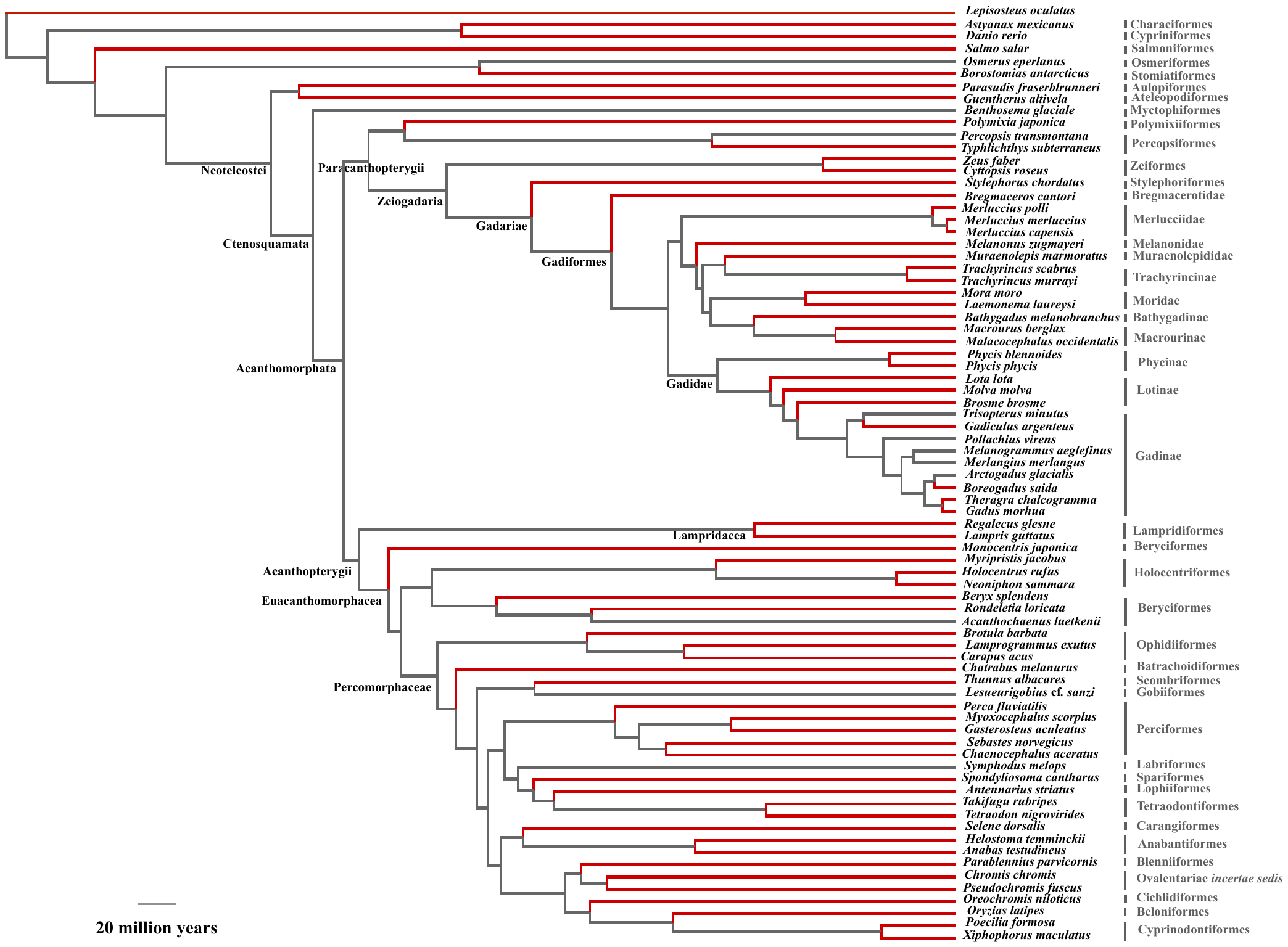

**Supplementary Figure 1 Sequence evolution of *lg*.** a. Comparative genomic analyses showed that *lg* is a fish-specific gene. b. Multiple losses occurred in different lineages within teleost fishes. The reference tree is from the literature ^1^.

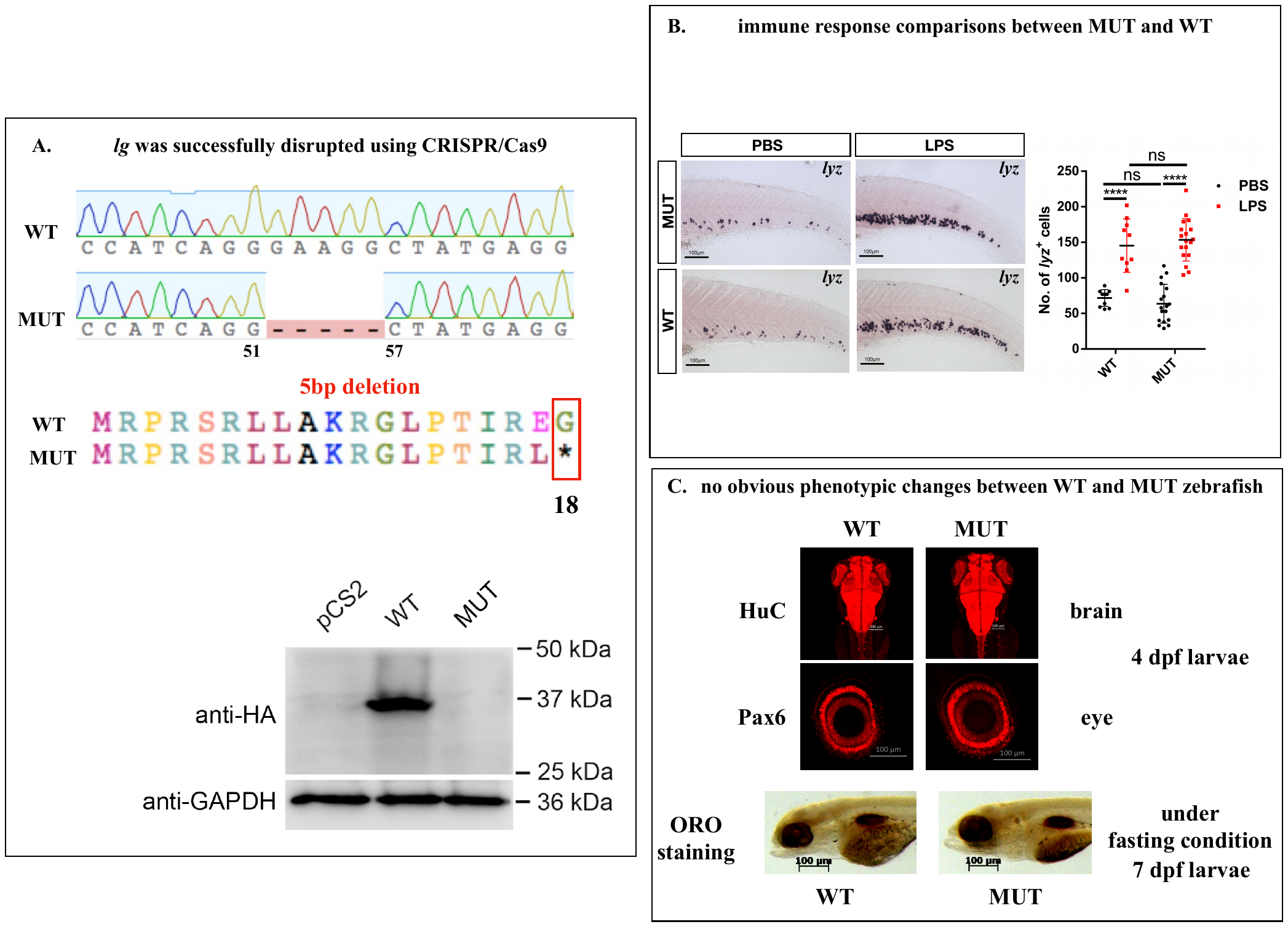

**Supplementary Figure 2 *lg* gene knockout in zebrafish and comparisons between mutant (MUT) and wild-type (WT) zebrafish.** A. *lg* was successfully disrupted using CRISPR/Cas9. This was confirmed by both Sanger sequencing (top) and western blots (bottom). B. Immune response comparisons between MUT and WT using whole mount *in situ* hybridization in CHT (caudal haematopoietic tissue) treated with LPS and PBS (n=10 for WT, and n=18 for MUT). Quantification of the numbers of *lyz*^+^ signals across the eight somites is shown as dots. n.s. not significant. Mean and SD error bars are shown. C. Phenotype comparisons between MUT and WT zebrafish. Brain and eye phenotypes in the mutants were examined with the neuron marker HuC. No obvious phenotype was found. Under fasting conditions, no obvious mutant phenotype was detected with ORO staining using the same strategy as described in the literature ^2^.

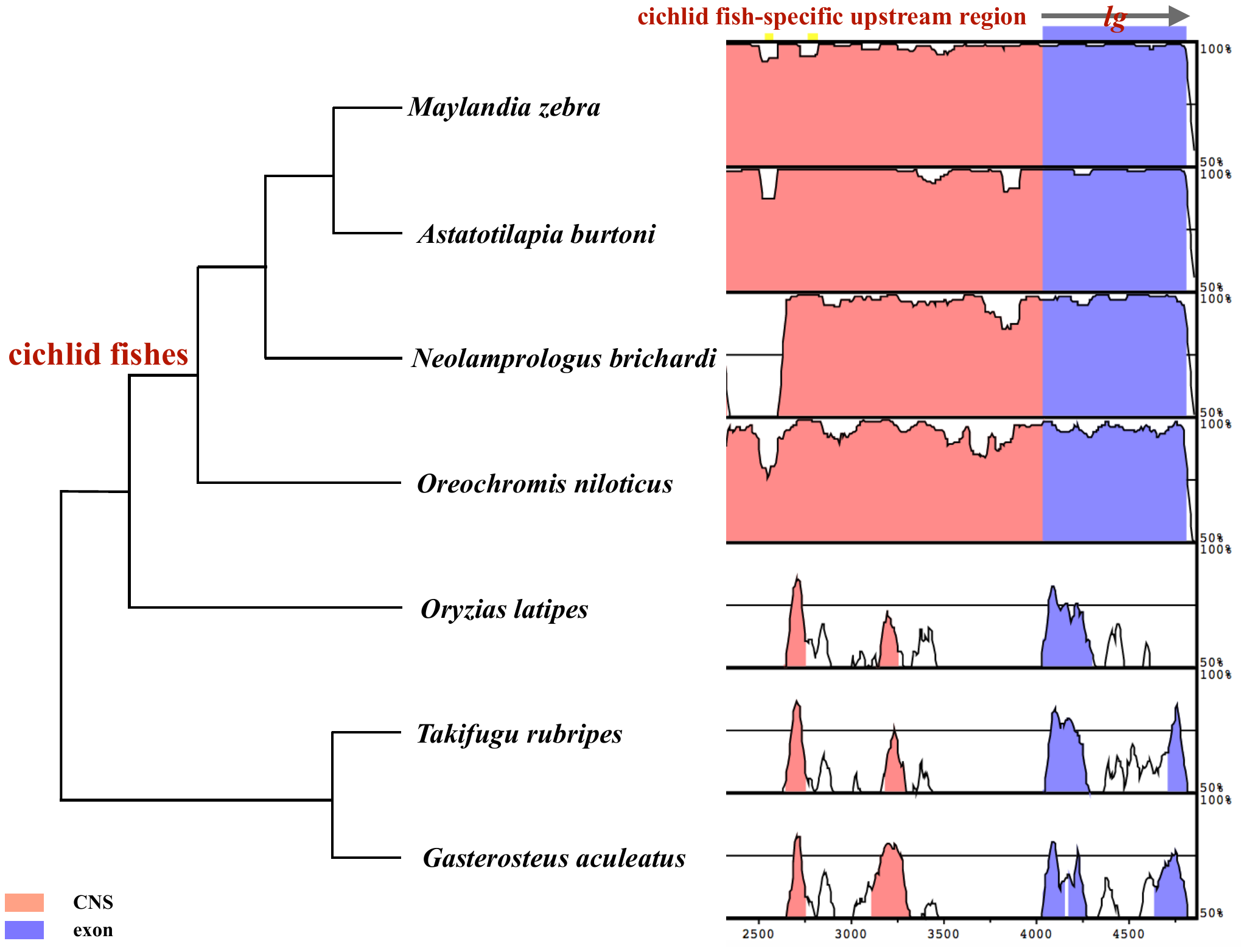

**Supplementary Figure 3** Genomic alignment of *lg* of cichlid fishes and other teleost fishes. mVISTA plots of a 2.5 kb region of the *lg* locus. The genome sequence of *Pundamilia nyererei* was used as the reference for the alignment. The pink regions are CNS (conserved noncoding sequences) regions. The dark blue regions are exons of *lg*. Cichlid fishes showed a large portion of cichlid fish-specific CNS in the upstream region.

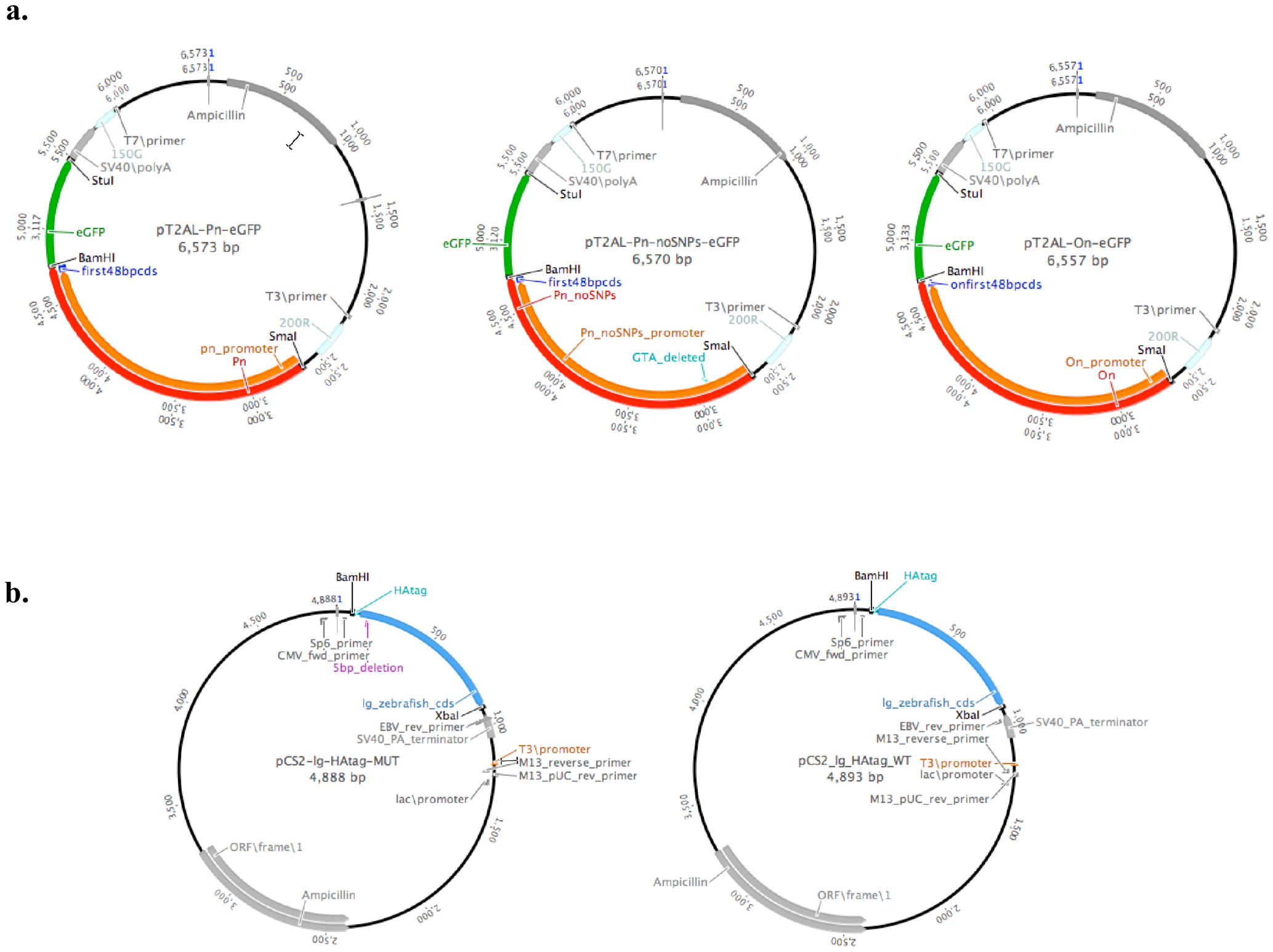

**Supplementary Figure 4** Schematic plasmid construction. (a) Different transgenic plasmids construction. Inserted cichlid fish sequences are shown in red, which include the target upstream region (shown in orange) and the first 48 bp coding region (shown in dark blue). To determine whether the cichlid fish-specific upstream region of *lg* has regulatory functions and whether this region of different cichlid fish lineages has different roles, we constructed different transgenic plasmids with the vector pT2AL to drive eGFP expression in zebrafish. The left plasmid has an upstream region of a modern haplochromine (MH) cichlid fish*, Pundamilia nyererei* (pT2AL-Pn-eGFP), as the promoter. The middle plasmid has the corresponding homologous region of a cichlid fish belonging to a sister lineage of East African Great Lakes cichlid fishes, Nile tilapia (*Oreochromis niloticus*) (pT2AL-On-eGFP). The right plasmid has the same upstream region as the Pn line but we deleted three MH-segregated SNPs (GTAs) (pT2AL-Pn-noSNPs-eGFP). (b) HAtag plasmid construction for western blot analyses between mutant (MUT) and wild-type (WT) zebrafish. Since there is no available antibody for *lg*, we thus added an HA tag (light green) in front of the coding region of *lg* (light blue) by PCR amplification*.* To ligate the amplicon of the HA-tagged coding region of MUT (left) and WT (right) zebrafish into the pCS2 vector, restriction enzyme sites (BamHI and XbaI) were added to the two ends of the amplicons by PCR amplification, followed by double digestion (NEB) and ligation (Promega). The antibody of the HA tag can thus detect the fused *lg* protein for the following western blot experiment.
